# Base-resolution prediction of transcription factor binding signals by a deep learning framework

**DOI:** 10.1101/2021.11.01.466840

**Authors:** Qinhu Zhang, Ying He, Siguo Wang, Zhanheng Chen, Zhenhao Guo, Zhen Cui, De-Shuang Huang

## Abstract

Transcription factors (TFs) play an important role in regulating gene expression, thus the identification of the sites bound by them has become a fundamental step for molecular and cellular biology. In this paper, we developed a deep learning framework leveraging existing fully convolutional neural networks (FCN) to predict TF-DNA binding signals at the base-resolution level, called FCNsignal. The proposed FCNsignal can simultaneously achieve the following tasks: (i) modeling the base-resolution signals of binding regions; (ii) discriminating binding or non-binding regions; (iii) locating TF-DNA binding regions; (iv) predicting binding motifs. The experimental results on 53 TF ChIP-seq datasets and 6 chromatin accessibility ATAC-seq datasets show that our proposed framework outperforms some existing state-of-the-art methods. In addition, we explored to use the trained FCNsignal to locate all potential TF-DNA binding regions on a whole chromosome and predict DNA sequences of arbitrary length, and the results show that our framework can find most of the known binding regions and accept sequences of arbitrary length. Furthermore, we demonstrated the potential ability of our framework in discovering causal disease-associated single-nucleotide polymorphisms (SNPs) through a series of experiments.

**Author summary:** Identification of transcription factor binding sites (TFBSs) is fundamental to study gene regulatory networks in biological systems, as TFs activate or suppress the transcription of genes by binding to specific TFBSs. With the development of high-throughput sequencing technologies and deep learning (DL), several DL-based approaches have been developed for systematically studying TFBSs, achieving impressive performance. Nevertheless, these methods either excessively focus on discriminating binding or non-binding sequences or individually accomplish multiple TFBSs-associated tasks. In this work, we provide an integrated framework, which utilizes the FCN architecture to predict TF-DNA binding signals at the base-resolution level, to simultaneously study multiple TFBSs-associated tasks. More importantly, we also demonstrate that our proposed framework has the ability to locate all potential TF-DNA binding regions from DNA sequences of arbitrary length. We hope that our framework can provide a new perspective on studying the mechanism of TF-DNA binding and its related tasks.

## Introduction

Transcription factors (TFs) can activate or suppress the transcription of genes by binding to specific DNA non-coding regions, thereby playing an integral role in gene expression [1]. These specific regions bound by TFs are known as transcription factor binding sites (TFBSs) [2], and the aligned profiles of TFBSs are referred to as *cis*-regulatory motifs [3]. In recent years, a lot of substantial computational efforts have been invested to study TF-DNA binding specificities and motifs prediction, resulting in the development of numerous algorithms, computational tools, and databases [4-7]. However, the deep understanding of the mechanism of TF-DNA binding remains fragmented. Thus, comprehensive computational methods are needed to systematically uncover the binding mechanism.

The fast development of high-throughput sequencing technologies has brought about a large amount of TF-DNA binding data for studying TFBSs-associated tasks. For example, ChIP-seq [8] provides an opportunity for viewing genome-wide interactions between DNA and specific TFs. Protein binding microarrays (PBMs) [9] have enabled large-scale characterization of TF-DNA binding in a high-throughput manner without considering the influence of cofactors on predicting TFBSs. ATAC-seq [10] describes putative accessible regions in the genome that often work together with transcription factors (TFs), RNA polymerases, or other cellular machines. Furthermore, SMiLE-seq [11] is a newly-developed technology for protein–DNA interaction characterization that can efficiently characterize DNA binding specificities of TF monomers, homodimers and heterodimers. These binding data provide an unprecedented opportunity for us to develop computational approaches to predict TFBSs and motifs. For example, MEME [12] discovered TF-DNA binding motifs by searching for repeated, ungapped sequence patterns that occur in the biological sequences. STREME [13] identified ungapped motifs with recurring, fixed-length patterns that are enriched in query sequences or relatively enriched in them compared to control sequences. gkm-SVM [14,15] detected functional regulatory elements in DNA sequences by using gapped kmer and support vector machine. However, these methods are often subject to the defects of low efficiency and poor performance.

There is hence a critical need for improved computational methods that can accurately identify cis-regulatory motifs on high-throughput sequencing data. In recent years, deep learning (DL) has achieved amazing performance in many fields, such as computer version [16,17] and natural language processing [18,19], inspiring researchers to develop DL-based methods for predicting TFBSs and motifs [20-26]. For example, DeepBind [27] and DeepSea [28] applied convolutional neural networks (CNN) to accurately predict diverse molecular phenotypes, including TF binding from DNA sequences. DanQ [29] predicted TF-DNA binding motifs and prioritized functional SNPs by combining CNN with recurrent neural network (RNN). However, these DL-based methods mainly focus on discriminating binding or non-binding sequences and fail to accurately predict TFBSs and motifs. To remedy such problems, DESSO [30] used a CNN model for extracting motif patterns from given ChIP-seq peaks and a statistical model based on the binomial distribution for optimizing the identification of motif instances. FCNA^*^ [31] predicted TFBSs and motifs on ChIP-seq data by using a fully convolutional neural network (FCN) and global average pooling (GAV). D-AEDNet [32] adopted an encoder-decoder architecture to identify the location of TF-DNA binding sites in DNA sequences. BPNet [33] introduced a dilated CNN to predict base-resolution ChIP-nexus binding profiles of pluripotency TFs. Although these DL-based methods have achieved high accuracy in the tasks of classification, location, or predicting motifs, a novel method that can integrate the above TFBSs-associated tasks in a single-task way is still missing.

Overall, we proposed a FCN-based framework (FCNsignal) to predict transcription factor binding signals at the base-resolution level, which can simultaneously achieve the following tasks: (i) modeling the base-resolution signals of binding regions; (ii) discriminating binding or non-binding regions; (iii) locating TF-DNA binding regions; (iv) predicting binding motifs. Specifically, the peaks and signals (*p*-value) of ChIP-seq and ATAC-seq were firstly collected from ENCODE. Secondly, FCNsignal took as input the peaks and predicted the base-resolution signals. Finally, several competing methods were used to evaluate the performance of FCNsignal on the four tasks. Experimental results show that FCNsignal is superior to the competing methods on these tasks, which demonstrates the effectiveness of our proposed method. In addition, we explored to use the trained FCNsignal to predict DNA sequences of arbitrary length and locate all potential TF-DNA binding regions on a whole chromosome, and the results show that our framework can accept inputs of arbitrary length and find most of the known binding regions. Furthermore, we demonstrated the potential ability of our framework in discovering causal disease-associated single-nucleotide polymorphisms (SNPs) through a series of experiments. Thus, we hope that FCNsignal can provide a new perspective on studying TFBSs and its related tasks.

## Results

### Overview of experimental design

FCNsignal, consisting of an encoder architecture, a decoder architecture, and a skip architecture, is designed to predict base-resolution signals of TF-DNA binding (Fig 1A). Substantially, FCNsignal is a base-resolution regression framework, which accepts DNA sequences where each sequence is converted to a matrix in one-hot format with A = [1, 0, 0, 0], C = [0, 1, 0, 0], G = [0, 0, 1, 0], and T = [0, 0, 0, 1], and uses base-resolution binding signals (*p*-value) as the supervised labels. Nevertheless, FCNsignal can also be applied to discriminate binding or non-binding regions, locate TF-DNA binding regions and predict binding motifs by using the maximum values of binding signals (see **Materials and methods**). A total of 53 ChIP-seq and 6 ATAC-seq datasets were used to build models and evaluate the predictive performance of FCNsignal.

**Fig 1.**
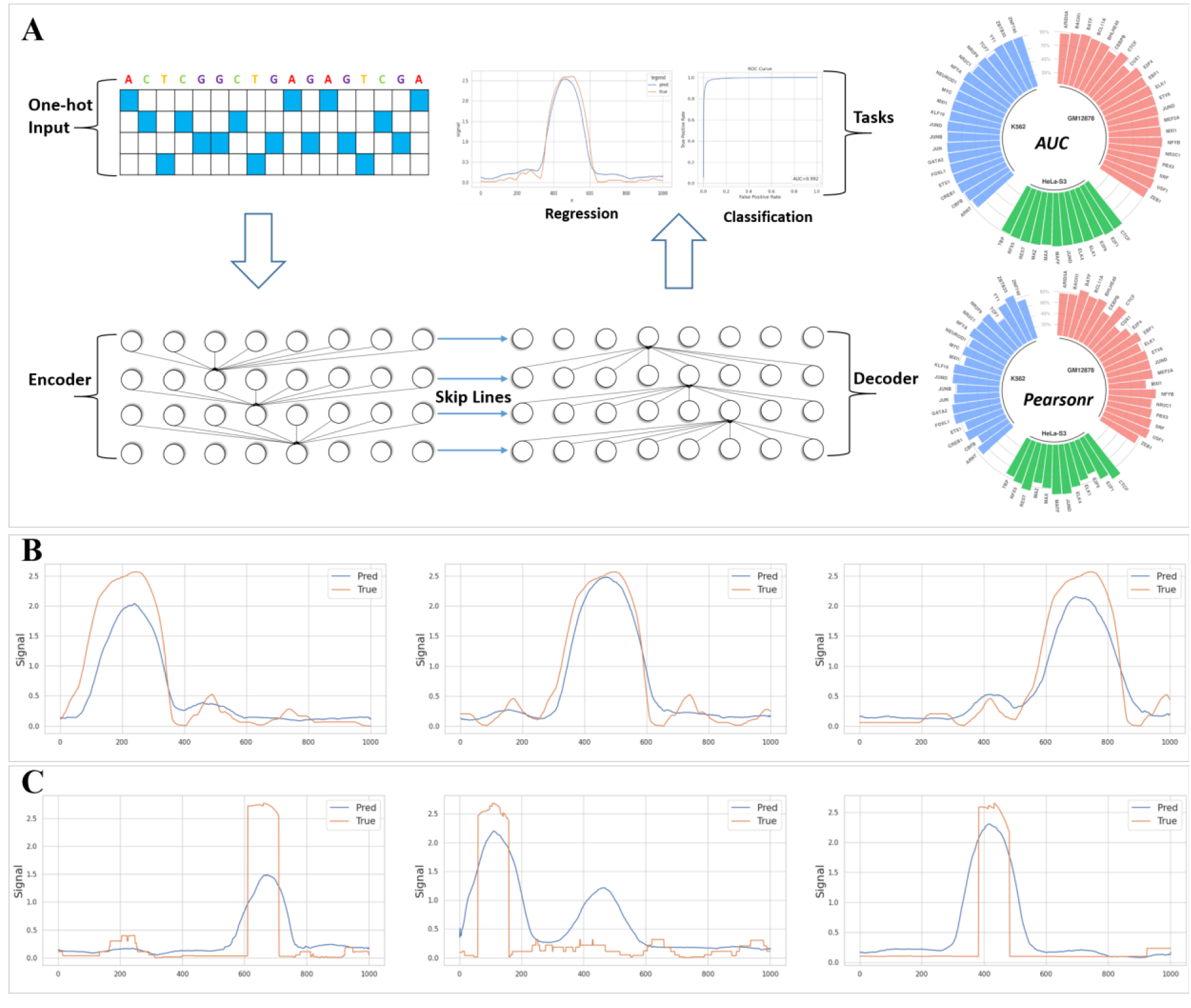
Schematic overview of FCNsignal framework. (A) FCNsignal is mainly composed of an encoder architecture, a decoder architecture, and a skip architecture, which takes as input DNA sequences and predicts the base-resolution signals. FCNsignal can simultaneously realize multiple tasks with high accuracy by using the maximum values of predicted signals. (B) FCNsignal can accurately capture the signals of the shifted binding regions that are separately located at 250bp, 500bp, and 750bp. (B) FCNsignal can accurately capture the signals of the binding regions that were randomly inserted into negative sequences.

### FCNsignal can accurately capture TF-DNA binding patterns

To test the ability to capture TF-DNA binding patterns, two validation experiments were designed. Specifically, for the first experiment: (i) the maximum values of signals for peaks were located; (ii) the length of the peaks were expanded to 1000bp by centering on the position of the maximum values; (iii) the same process was repeated twice after shifting the position of the maximum values by -250bp and +250bp respectively. For the second one: (i) the maximum values of signals for peaks were located; (ii) the regions of length 100bp surrounding the position of the maximum values were extracted; (iii) the extracted regions were randomly inserted into negative sequences and kept to 1000bp. Subsequently, the trained FCNsignal was used to predict the base-resolution signals of these data. As shown in Fig 1B and 1C (S2 Fig), we observe that our proposed method can accurately identify the sequence-specific patterns of TF-DNA binding regions and model their corresponding signals.

### The overall performance of FCNsignal on ChIP-seq datasets

To test the performance of FCNsignal on ChIP-seq datasets, we collected 53 ChIP-seq datasets for sequence-specific TFs from the ENCODE project, which consists of 21, 20, and 12 TF datasets in the GM12878, K562, and HeLa-S3 cell lines respectively. The overall performance of the proposed method was investigated on the test data.

The MSE and Pearsonr between the predicted signals and the true signals were employed to evaluate the performance of FCNAsignal in modeling the base-resolution signals. FCNA and BPNet were used as the competing methods. As shown in Fig 2A, the average MSE values of FCNsignal are lower than that of other methods, and the average Pearsonr values are higher than that of other methods. More comparisons (S3A Fig) show that our method outperforms the other two methods under the MSE and Pearsonr metrics. These results indicate that FCNsignal can model the base-resolution signals well.

**Fig 2.**
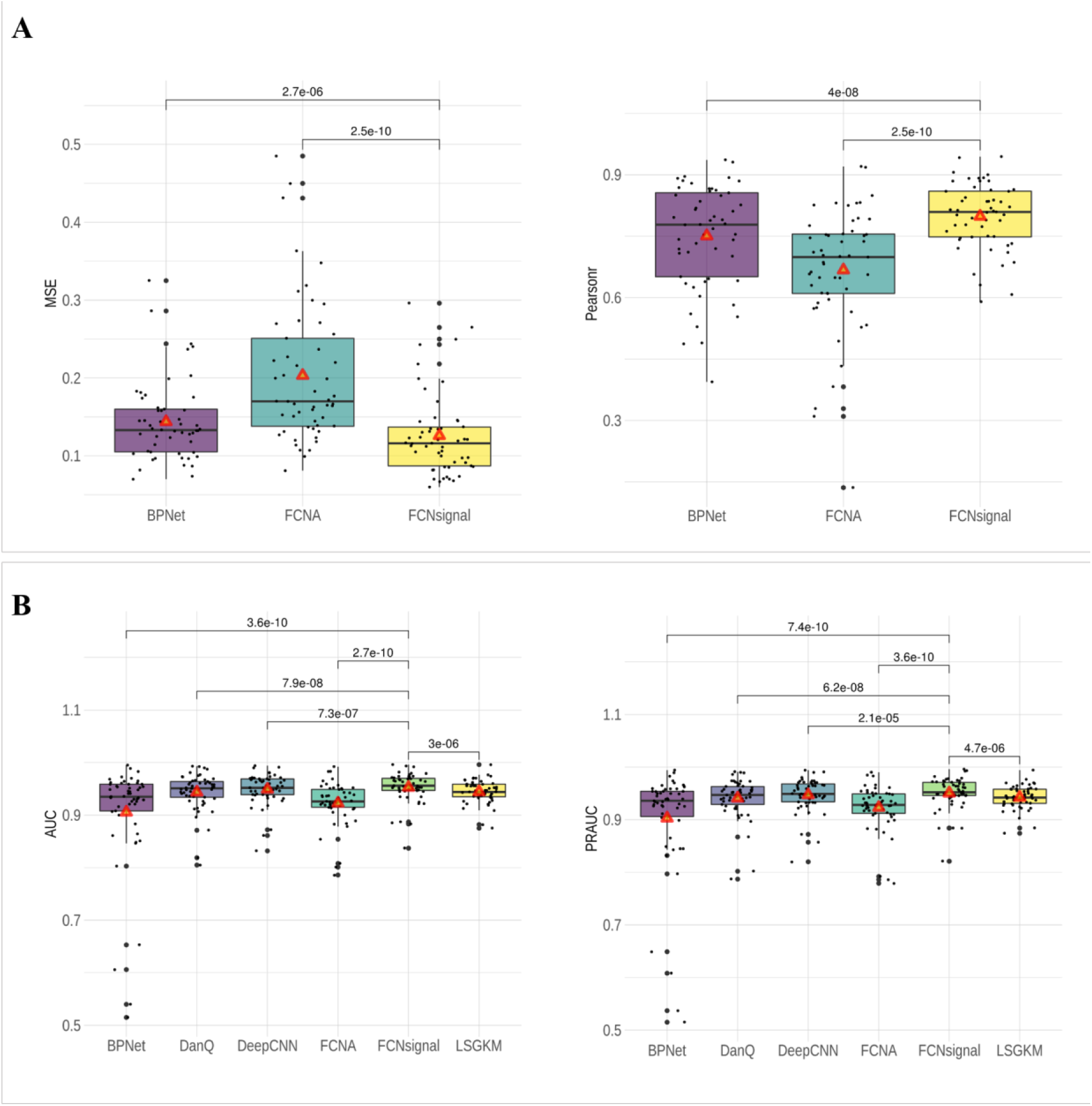
Performance comparison of FCNsignal and the competing methods on the 53 ChIP-seq datasets. (A) The MSE and Pearsonr values for BPNet, FCNA, and FCNsignal, and the Wilson’s test *p*-values (paired) of the two metrics between FCNsignal and the other two methods. (B) The AUC and PRAUC values for BPNet, DanQ, DeepCNN, FCNA, FCNsignal, and LSGKM, and the Wilson’s test *p*-values (paired) of the two metrics between FCNsignal and the other five methods. The red triangles represent the average values.

The AUC and PRAUC metrics were employed to evaluate the performance of FCNsignal in discriminating binding or non-binding sequences. LSGKM, DanQ, DeepCNN, FCNA, and BPNet were used as the competing methods. As shown in Fig 2B, our method outperforms the five competing methods with low p-values (Wilson’s test, paired), and the average AUC (PRAUC) values are higher than theirs. More comparisons are displayed in S3B Fig, but we observe that FCNsignal performs much worse than LSGKM on three TFs, including CEBPB (AUC: 0.883 vs. 0.913), CUX1 (AUC: 0.837 vs. 0.882), and TCF7 (AUC: 0.885 vs. 0.944). To analyze this phenomenon, for each of them, the distribution of the maximum values of signals in the positive and negative samples was calculated. As shown in S4 Fig, for each of the three TFs, the majority of the maximum values in the positive and negative samples are intersected so that it is difficult for our method to accurately discriminate positive or negative samples by the maximum values of signals. On the contrary, for CTCF, the maximum values in the positive and negative samples are barely intersected so that our method can accurately discriminate the positive or negative samples with superior performance over LSGKM (AUC: 0.993 vs. 0.955). Overall, these observations demonstrate that our method can accurately discriminate binding or non-binding sequences, but also is possibly influenced by the quality of signals.

To explore the effect of the number of samples on the performance of FCNsignal, three methods were selected including LSGKM, FCNAsignal, and BPNet, whose model complexity is increasing. All TF datasets were first sorted in ascending order of quantity, and then the three methods were compared using the AUC metric. As shown in S5 Fig, we observe that (i) LSGKM has the competitive performance to and even surpasses FCNsignal on the datasets with small size, but performs worse than FCNsignal on the datasets with large size; (ii) BPNet performs much worse than the other two methods on the datasets with small size but is getting better results as the number of datasets increases. Generally, these observations illustrate that FCNsignal is slightly affected by the number of samples but also benefits from the increasing number of samples.

### FCNsignal can accurately predict TF-DNA binding motifs

Seven competing methods, including MEME, STREME, DanQ, DeepCNN, FCNA^*^, FCNA, and BPNet, were employed to investigate the performance of FCNsignal in predicting TF-DNA binding motifs. As described in ‘**Materials and methods**’ section, the –log2(*p*-value) produced by TOMTOM was used as the evaluation metric. As shown in Fig 3A, we observe that (i) FCNsingal outperforms MEME, STREME, DanQ, DeepCNN, and BPNet with low *p*-values (Wilson’s test, paired), and the average –log2(p-value) values are higher than theirs; (ii) FCNA has the competitive performance to FCNsignal, but the average –log2(p-value) value is lower than FCNsignal; (iii) FCNA^*^ perform significantly better than FCNsignal, and the average –log2(p-value) values are the highest among all methods; (iv) the number of motifs found by FCNsignal (50/53) is more than other competing methods except FCNA^*^ that finds all motifs. FCNA and FCNsignal are both based on the symmetrical FCN architecture, so they have comparable performance. FCNA^*^ used the base-resolution labels (0/1) which were manually annotated by PCMs while FCNsignal used the base-resolution signals which were produced by high-throughput sequencing technologies, so FCNA^*^ achieves the very impressive performance. However, FCNA^*^ was extremely dependent on the PCMs so that it cannot distinguish positive sequences from negative sequences, which means that it even using the negative sequences can also get very good results (experimental details and results can be found in S1 Text and S6 Fig respectively), whereas FCNsignal can distinguish positive sequences from negative sequences by the maximum values of signals. Moreover, the real PCMs are not easily acquired unlike the signals produced by sequencing technologies. Generally, these results indicate that FCNsignal can accurately and efficiently predict TF-DNA binding motifs. Motif visualization of some examples is displayed in S3 Table.

**Fig 3.**
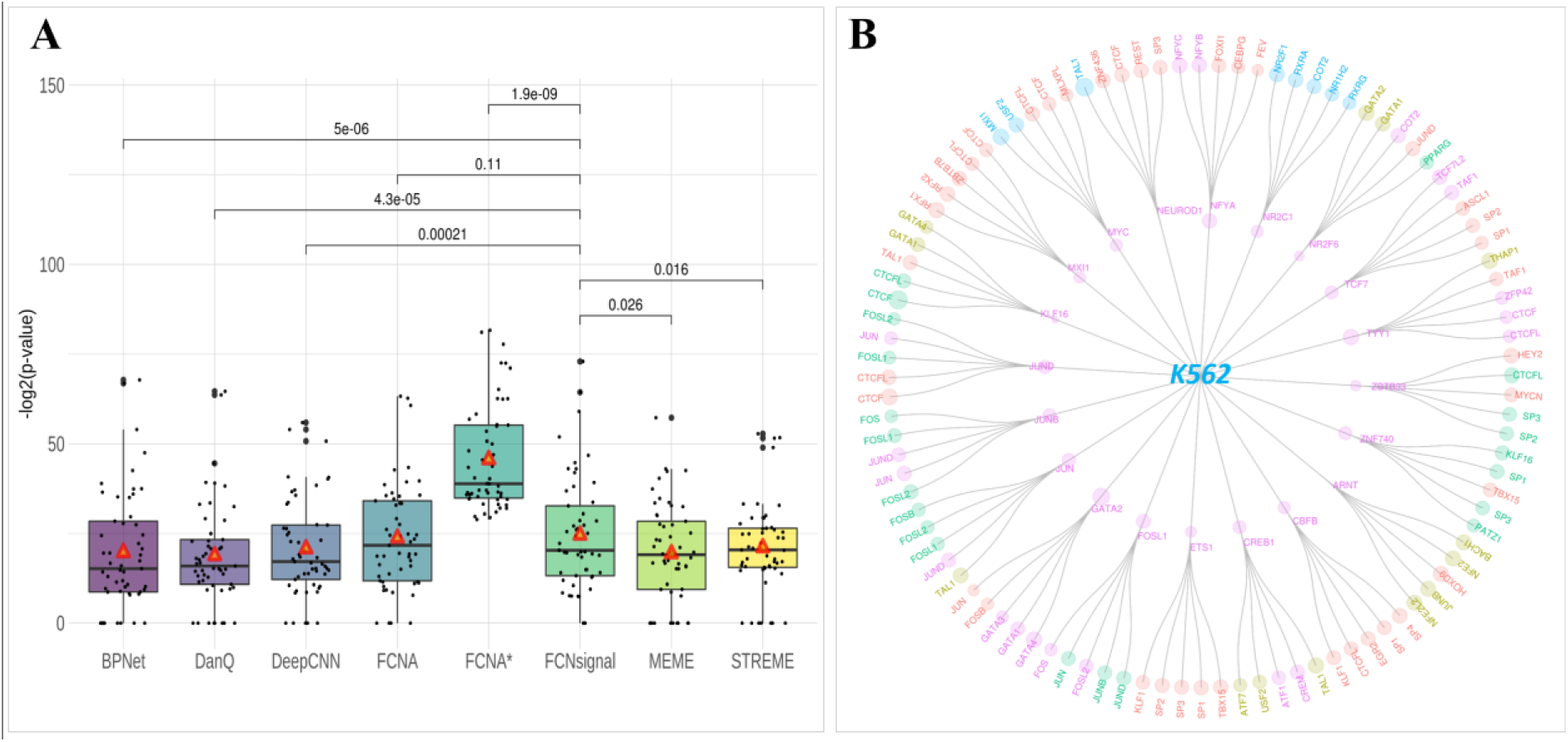
Motif prediction performance comparison of FCNsignal and the competing methods. (A) The –log2(*p*-value) values for BPNet, DanQ, DeepCNN, FCNA, FCNA^*^, FCNsignal, MEME and STREME, and the Wilson’s test *p*-values (paired) between FCNsignal and other seven methods. Note that the red triangles represent the average values and the value of 0 means not finding the target. (B) Detailed view of identified TF motifs for K562. The TF motifs of the inner loop are the target ones and the motifs of the outer loop are the found ones. The size of circles corresponds to the –log2(*p*-value) value and different colors are used to designate different TF classes. The TF motifs marked by ‘pink’ belong to the same TF family sharing the consensus binding sequence while the ones marked by other colors are more likely the indirect TF motifs.

In addition, FCNsignal can be used to find indirect TF-DNA binding motifs. Taking all TFs in the K562 cell line as an example, except each target TF motif, the top-5 matched motifs were picked out according to the – log2(p-value). As shown in Fig 3B, we observe that (i) FCNsignal are prone to finding similar motifs belonging to the same TF family. For example, the matched TF motifs of NFYA contain NFYB and NFYC, which belong to the Heteromeric CCAAT-binding family. The matched TF motifs of GATA2 contain GATA1, GATA3, and GATA4, which belong to the C4-GATA-related family; (ii) FCNsignal can find the indirect TF motifs interacting with the target TF. For instance, the Jun-related motifs (JUNB and JUND) and the Fos-related motifs (FOSL1 and FOSL2) are mutually inclusive, which has been demonstrated that all JUN–FOS heterodimers often strongly bind to the TPA-response element [11]. The matched TF motifs of GATA2 involve JUN and FOSB with significant p-values, which has been proved that GATA2 frequently occupies the same chromatin sites as c-JUN and c-FOS, heterodimeric components of AP-1 [34].

### FCNsignal can locate potential TF-DNA binding sites on the whole chromosome

At first, the cross-cell-line prediction ability of FCNsignal was tested by using the models trained on the datasets in the GM12878 and K562 cell lines to predict the datasets in the HeLa-S3 cell line, and these datasets include CTCF, ELK1, and JUND. As shown in S7A Fig, the performance of FCNsignal in predicting the datasets from different cell lines is slightly worse than it predicting the ones from the same cell lines, but the overall cross-cell-line prediction performance is very high. The above observations imply that FCNsignal is capable of predicting specific TF-DNA binding sites across the genome.

To test the ability of FCNsignal in predicting potential TF-DNA binding sites on the whole chromosome, we applied a chromosomally-split strategy to divide the experimental datasets into training data, validation data, and test data. Specifically, (i) six TF ChIP-seq datasets were picked out from the HeLa-S3 and K562 cell lines respectively; (ii) for each dataset, the data from chromosomes 17 and 18 were separately used as the test and validation sets while the remaining data were used as the training set; (iii) these data were used to train and test FCNsignal. The results demonstrate that FCNsignal can achieve high prediction performance regardless of using the chromosomally-split or randomly-split strategies (S7B Fig). Subsequently, the entire chromosome 17 was used to validate the performance of our proposed method in locating TF-DNA binding regions. Briefly, (i) the entire chromosome 17 was segmented into sequences of length 1000bp; (ii) the trained models for CTCF and YY1 were used to predict the base-resolution binding signals of these sequences; (iii) several sequences with high confidence were selected only if the maximum values of the predicted signals exceed a manually set threshold (here we set it to 1.5); (iv) the regions of length 60bp surrounding the position of the maximum values were extracted. As a result, 2132 and 1060 potential regions containing TFBSs for CTCF and YY1 were discovered respectively.

To validate these located short regions, the number of them intersecting with the true peaks collected from ReMap2020 was counted by the rule that the intersection ratio is over 50%. As shown in Fig 4A, 81% (1725/2132) of CTCF and 63% (671/1060) of YY1 are supported by the true peaks. Moreover, we analyzed the distribution of h3k27ac signals for supported and unsupported regions, finding that (i) the h3k27ac values of the supported regions are significantly higher than that of the unsupported regions, meaning that the supported regions are mostly from open chromatin regions while the unsupported regions are perhaps from unopen regions, and (ii) the unsupported regions also contain a small number of outliers that have high h3k27ac values, implying that these outliers are similar to the CTCF/YY1-associated binding sites. Besides, we utilized FIMO [35] to find motif instances with high significance (*p*-value < 1e-4) and CentriMo [36] to do motif enrichment analysis, on the located regions and the negative regions respectively. As opposed to selecting the located regions, the sequences were chosen as the negative regions if the maximum values of the predicted signals are less than the threshold, of which the same number as the located ones were randomly selected. As a result, the number of motif instances discovered by FIMO on the located regions (1682 for CTCF and 514 for YY1) is much more than the ones found on the negative regions (117 for CTCF and 34 for YY1), and the –log2(*p*-value) values of the located regions are much higher than that of the negative regions (Fig 4B). Moreover, motifs for CTCF and YY1 are also more enriched on the located regions than that on the negative regions (Fig 4C). These results confirm that the located regions are more likely to contain TFBSs. More examples can be found in S7 Fig, which have the same observations. Overall, FCNsignal has the capability of classifying and locating potential TF-DNA binding sites on the whole chromosome but does not easily distinguish similar binding sites (e.g. bound by the TFs from the same family).

**Fig 4.**
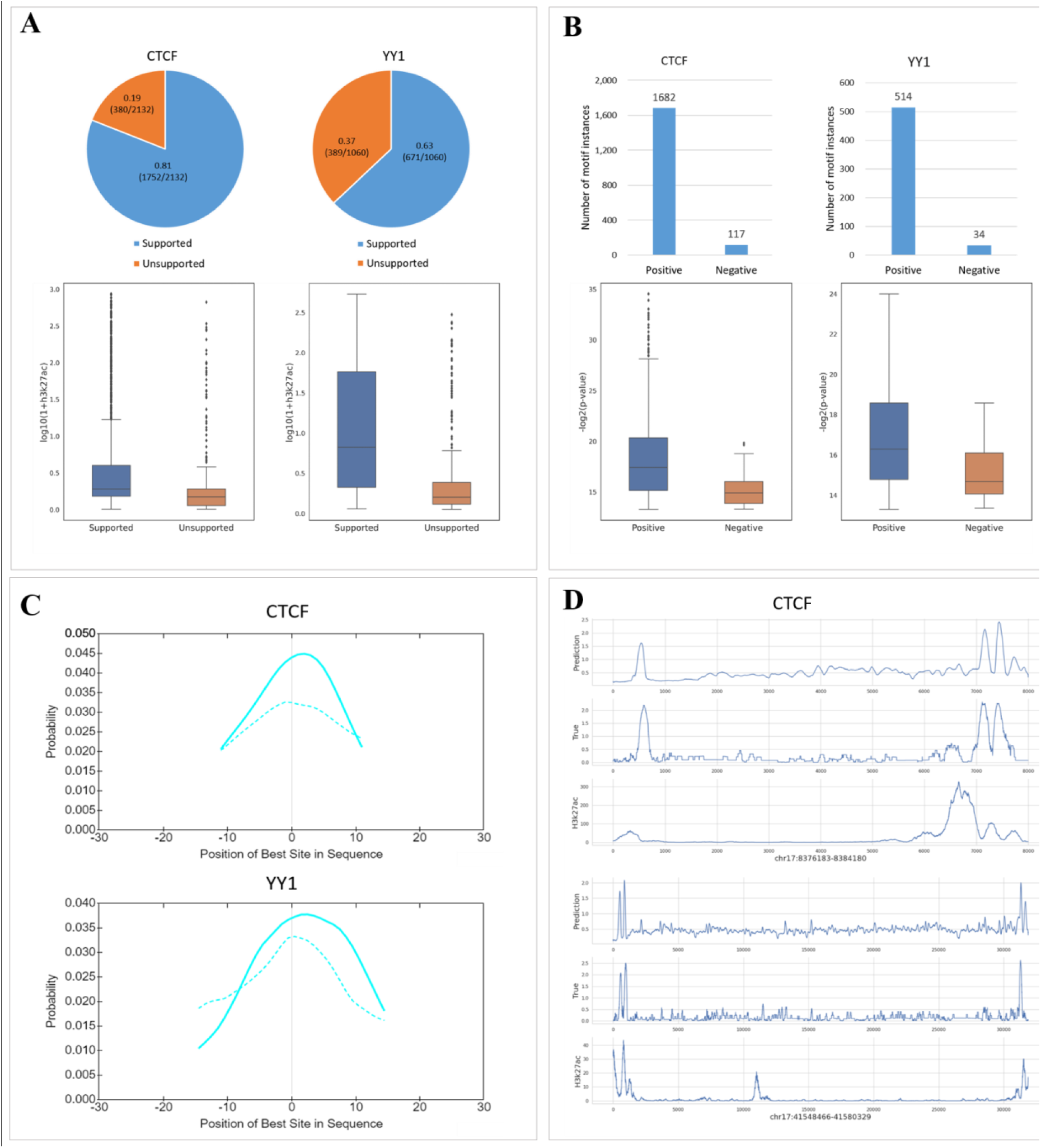
The performance of FCNsignal in locating TF-DNA binding regions. (A) The results of FCNsignal in locating potential binding regions for CTCF and YY1 on the whole chromosome 17 as well as the distribution of h3k27ac signals in the supported and unsupported sequences, where ‘Supported’ means existing in the real peaks and ‘Unsupported’ means the opposite. (B) The number of motif instances found by FIMO on the located regions and the negative regions as well as the distribution of the matched –log2(*p*-value) values. (C) The motif enrichment analysis on the located regions and the negative regions. The dash lines represent the results on the negative regions. (D) The base-resolution signals of DNA sequences of arbitrary length predicted by FCNsignal, the true signals, and the h3k27ac signals. The range of the top DNA sequence is chr17: 8376183-8384180 (7997bp length), and the range of the bottom DNA sequence is chr17: 41548466-41580329 (31863bp length).

Since FCNsignal is a FCN-based framework, so it can accept DNA sequences of arbitrary length. To explain this, we randomly cut off a few sequences of arbitrary length that contain multiple peaks from the chromosome 17. Then, the trained models for CTCF and YY1 were used to predict the base-resolution binding signals of these sequences. As shown in Fig 4D, FCNsignal can take sequences of arbitrary length as input and accurately predict the base-resolution binding signals. More instances can be found in S8 Fig.

### The overall performance of FCNsignal on ATAC-seq datasets

To test the performance of FCNsignal on ATAC-seq datasets, we collected 6 ATAC-seq datasets from the ENCODE project, including A549, GM12878, HepG2, IMR90, K562, and MCF7 cell lines. The way of processing ATAC-seq datasets is the same as ChIP-seq datasets. It is well known that ATAC-seq data are often applied to the task of predicting chromatin accessibility that is equivalent to identifying open or unopen regions across the genome. In this study, we applied FCNsignal to predicting chromatin accessibility. Briefly, FCNsignal accepted DNA sequences and predicted their corresponding base-resolution signals, and then used the maximum values of signals to distinguish open regions from unopen regions. Four competing methods including LSGKM, DeepEmbed, Deopen, and BPNet were used to compare with our proposed method.

The MSE and Pearsonr metrics were adopted to compare the performance of FCNsignal and BPNet in modeling the base-resolution signals. As shown in Fig 5A, FCNsignal performs better than BPNet on all datasets. The AUC and PRAUC metrics were employed to compare the performance of FCNsignal and the four competing methods in discriminating binding or non-binding sequences. As shown in Fig 5B, FCNsignal outperforms the four competing methods on all datasets except that BPNet is superior to FCNsignal on the A549 dataset. Moreover, BPNet has comparable performance to FCNsignal when the number of samples is abundant, which confirms the aforementioned conclusion that BPNet performs poorly on the datasets with small size but gets good results on the datasets with large size. Overall, the above observations demonstrate that the performance of FCNsignal on ATAC-seq datasets is better than other competing methods.

**Fig 5.**
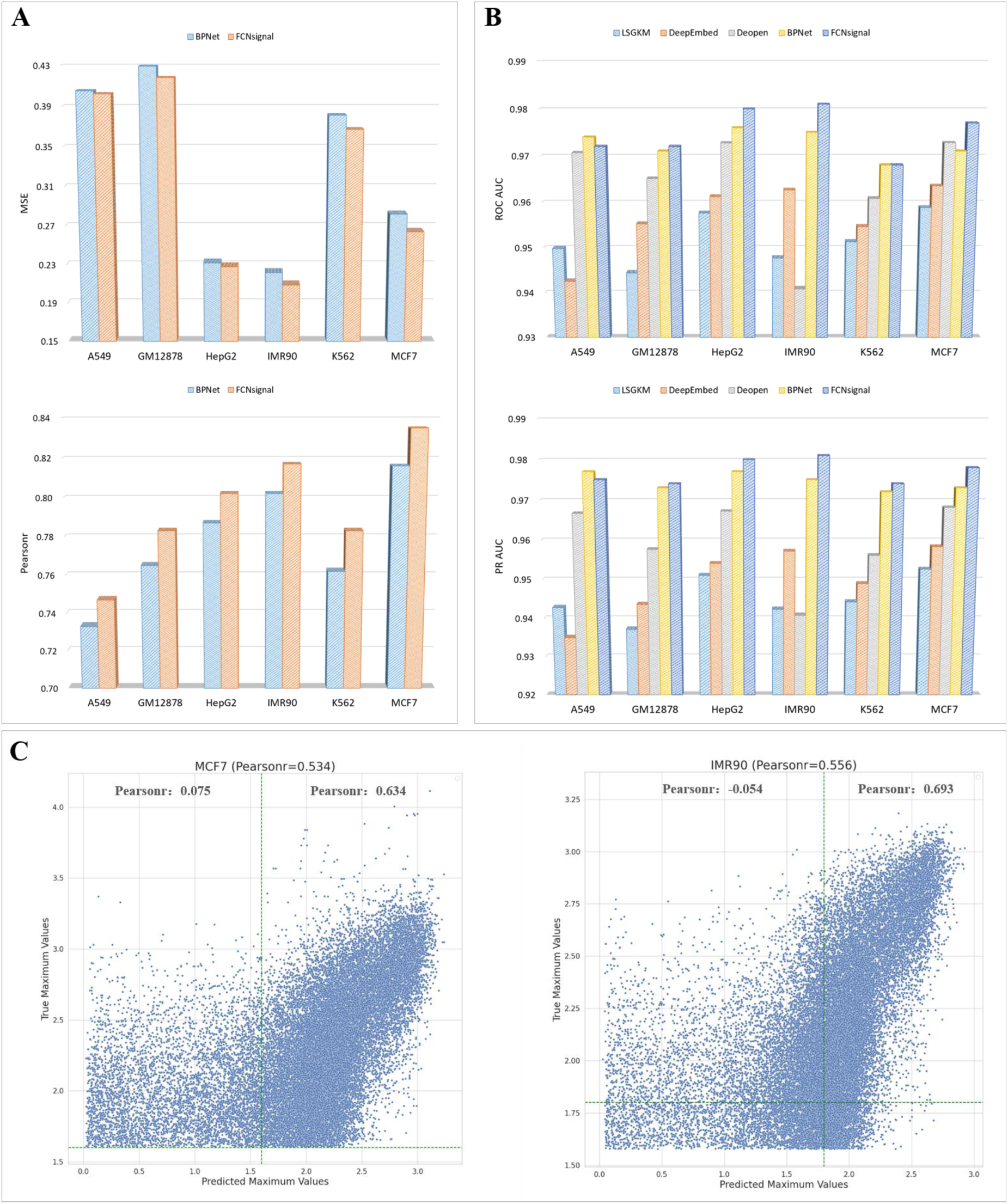
Performance comparison of FCNsignal and the competing methods on the six ATAC-seq datasets. (A) The MSE and Pearsonr values for BPNet and FCNsignal. (B) The AUC and PRAUC values for LSGKM, DeepEmbed, DeopenBPNet, and FCNsignal. (C) The Pearsonr between predicted maximum signals and true maximum signals for MCF7 (Pearsonr: 0534) and IMR90 (Pearsonr: 0.556). For MCF7, the Pearnsonr for DNA sequences with low and high openness are 0.075 and 0.634 respectively. For IMR90, the Pearnsonr for DNA sequences with low and high openness are -0.054 and 0.693 respectively.

In the above experiments, we simply considered the general regression and classification performance of FCNsignal. However, the degree of accessibility of DNA sequences may differ from each other even when they have the same binary labels. Such difference in the degree of accessibility implicates that classification models are unable to discriminate putative open regions with different accessibility. To explore this problem, we still used the maximum values of predicted signals to model the degree of accessibility of ATAC-seq peaks. Therefore, the openness of DNA sequences from the original test data, with only positive samples included, across different cell lines were predicted using the trained FCNsignal. As shown in Fig 5C and S9 Fig, we observe that FCNsignal performs well for DNA sequences with high openness values (mean Pearsonr: 0.62) but poorly for ones with low openness values (mean Pearsonr: 0.07). The possible reason leading to this phenomenon is that the peak signals are prone to noise, especially for those with weak binding signals. Nevertheless, the task of discriminating putative open regions with different accessibility remains a big challenge.

In addition, the enrichment of different TFBSs in each cell was analyzed by utilizing the trained FCNsignal. Briefly, (i) all matched TF motifs were found by following the process of motifs prediction (see **Materials and methods**); (ii) the significant matched motifs were picked out by filtering out those with *p*-value less than 1e-04; (iii) the number of occurrences of each motif was counted as the enrichment value. The heat maps of these selected TF motifs across the six cell lines are plotted in Fig 6A, obviously showing some TFBSs enrichments in the different cell lines. For example, (i) CTCF and Sp1-like family (e.g. Sp1, Sp2) motifs widely exist in the six cell lines; (ii) Ets-related family (e.g. ETV1, ETV6) motifs are enriched in the GM12878 cell line; (iii) AP2-related family (e.g. TFAP2A, TFAP2B) motifs are enriched in the MCF7 cell line; (iv) RXR-related family (e.g. NR2F1, NR2C2) motifs are enriched in the HepG2 cell line; (v) TEF1-related family (e.g. TEAD1, TEAD3) motifs are enriched in the IMR90 cell line. To explain this, we collected all non-redundant peaks of five TFs including CTCF, ETV6, TFAP2A, NR2C2, and TEAD1 from ReMap2020 and computed the intersection ratio between them and the ATAC-seq peaks of the six cell lines. As shown in Fig 6B, the intersection ratio of CTCF to the six cell lines is all very high, while the maximum intersection ratio of other TFs (ETV6, TFAP2A, NR2C2, and TEAD1) lies in the GM12878, MCF7, HepG2, and IMR90 cell lines respectively, which is consistent with the above observations (Fig 6A). To sum up, the motifs learned by FCNsignal can reflect the enrichment of TFBSs in the different cell lines.

**Fig 6.**
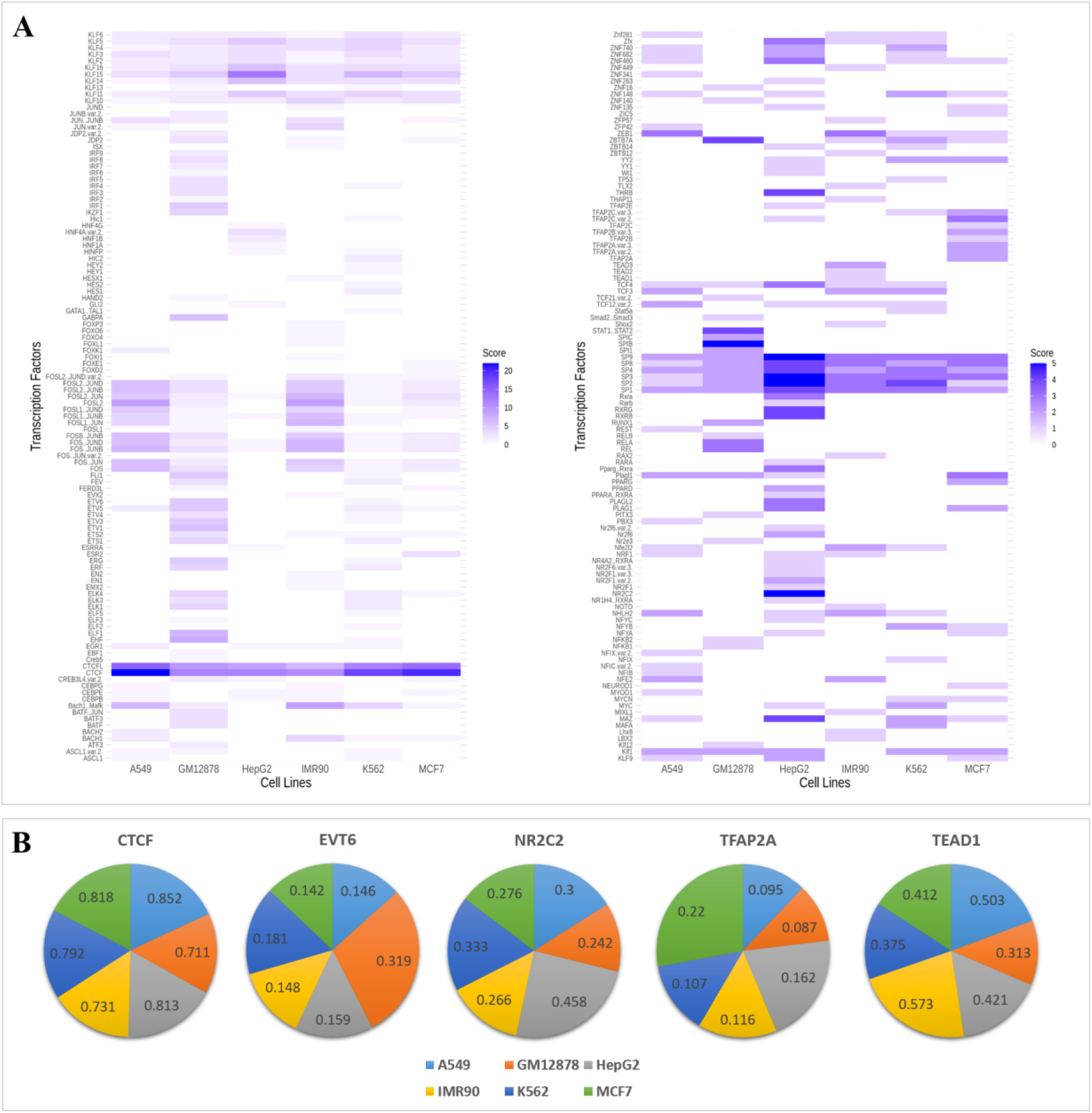
The enrichment of different TFBSs in the six cell lines. (A) The heat maps of diverse motifs in the six cell lines. CTCF binding motif is enriched in all six cell lines. EVT6, NR2C2, TFAP2A, and TEAD1 binding motifs are enriched in the GM12878, HepG2, MCF7, and IMR90 respectively. (B) The intersection ratio between the above five TFs ChIP-seq peaks and the ATAC-seq peaks of the six cell lines. The intersection ratio of CTCF to the six cell lines is all very high (mean: 0.786) corresponding to the observation that CTCF binding motif is enriched in all six cell lines. The maximum intersection ratio of the other four TFs is 0.319, 0.458, 0.22, and 0.573, corresponding to the GM12878, HepG2, MCF7, and IMR90 cell lines respectively.

### FCNsignal can identify causal SNPs from LD groups

To test the ability of FCNsignal in identifying causal SNPs, four causal SNP datasets with strong LD effects were used and DeltaSVM was chosen as the competing method. For myeloma, pan-autoimmune, and CLL, the FCNsignal trained on the GM12878 cell line was used to compute the SNP scores. For cancer breast, the FCNsignal trained on the MCF7 cell line was used to compute the SNP scores (see **Materials and methods**). DeltaSVM computed the SNP scores by directly utilizing the outputs of the LSGKM models on the GM12878 and MCF7 cell lines. Subsequently, these SNPs are prioritized by the SNP scores and the highest scored SNP in each set is assumed to be the causal one. As shown in Fig 7A, both FCNsignal and DeltaSVM successfully identify the causal SNPs in myeloma risk variants (rs4487645) and CLL risk variants (rs539846). Nevertheless, DeltaSVM fails to correctly identify the causal SNP (rs6927172) among the seven pan-autoimmune genetic susceptibility candidate SNPs while FCNsignal is able to pinpoint the right one. For breast cancer variants, Fig 7B shows that the distribution of the SNP scores of LD groups predicted by FCNsignal is more concentrated in the low score region than the ones predicted by DeltaSVM. Moreover, rs4784227 is believed to disrupt the binding of FOXA1 [37], and rs3803662 is in significant LD with it in individuals of European ancestry. As a result, the ratio of their SNP scores predicted by FCNsignal (0.054/0.001=54) is much higher than that predicted by DeltaSVM (2.28/1.54=1.5). The above results demonstrate that our proposed method can identify causal SNPs from LD groups. Besides, BPNet is also able to successfully identify causal SNPs from LD groups (S10 Fig), which demonstrates that the frameworks for predicting base-resolution signals can be directly used to identify causal SNPs.

**Fig 7.**
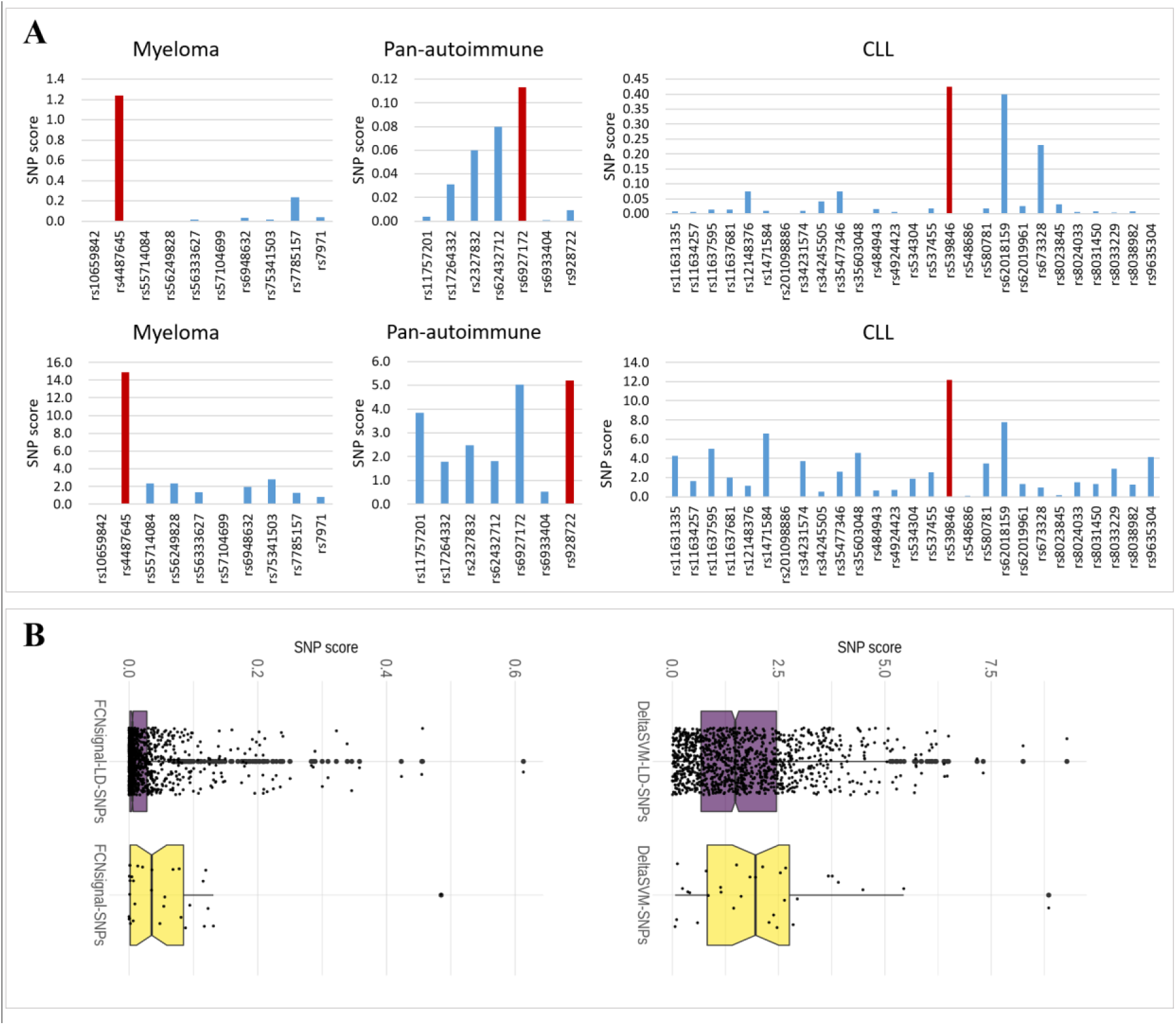
FCNsignal successfully pinpoints causal disease-related SNPs from LD groups. (A) The SNP scores of FCNsignal (top) and DeltaSVM (bottom) on the myeloma, pan-autoimmune, and CLL. The risk variants for the myeloma, pan-autoimmune, and CLL are rs4487645, rs6927172, and rs539846 respectively. (B) The distribution of the SNP scores of cancer breast predicted by FCNsignal (left) and DeltaSVM (right). The SNP scores of LD groups predicted by FCNsignal are more concentrated in the low score region than the ones predicted by DeltaSVM.

## Discussion

In this paper, we proposed a novel FCN-based framework, named FCNsignal, to predict transcription factor binding signals at the base-resolution level, followed by its application on the human ChIP-seq datasets and ATAC-seq datasets collected from ENCODE. This work provides an integrated framework to achieve multiple TFBSs-related tasks, including (i) modeling the base-resolution signals of binding regions; (ii) discriminating binding or non-binding regions; (iii) locating TF-DNA binding regions; (iv) predicting binding motifs, while amounts of previous works are always individually concentrated on these tasks and rarely on the task of locating binding regions. We validated that the maximum values of the base-resolution signals can reflect the openness degree of TF-DNA binding. In view of this, the maximum values can be employed to discriminate binding or non-binding regions as the openness degree of binding regions is far higher than that of non-binding regions, and also used to locate binding regions as the maximum values are most likely located in the binding regions. Our experimental results on the ChIP-seq and ATAC-seq datasets show that FCNsignal outperforms several existing state-of-the-art methods in the tasks of signals regression, sequences classification, and motifs prediction. Additionally, FCNsignal has the ability to classify and locate potential binding regions on the whole chromosome (e.g. the chromosome 17) and can predict the base-resolution signals of DNA sequences of arbitrary length. Moreover, FCNsignal trained on the ATAC-seq datasets can be directly used to identify causal SNPs from LD groups.

Although FCNsignal achieves superior performance in the four tasks, it still has several limitations: (i) FCNsignal relies on one-hot encoding, which suffers from an obvious limitation of being unable to capture dependencies between nucleotides. Hence, integrating *k*-mer based approaches [38,39] into FCNsignal would be considered; (ii) FCNsignal for modeling the degree of accessibility of ATAC-seq peaks is susceptible to noise, especially for the regions with weak binding signals. Hence, quality control of datasets is very essential for FCNsignal; (iii) FCNsignal does not easily discriminate similar binding sites (e.g. bound by the TFs from the same family), which is a common problem that almost all DL-based frameworks will encounter; (iv) The way of generating negative sequences is relatively simple (e.g. the upstream or downstream of peaks). Hence, the complex negative sequences should be considered (e.g. similar to the positive sequences or matched GC content with the positive sequences [40]); (v) Model interpretation of FCNsignal is weak, thus some advanced interpretation techniques, such as Grad-CAM [41], DeepLIFT [42], would be applied to extract more complex rules for TF-DNA binding.

Except for the above limitations, some interesting problems should also be explored in future works. For example, (i) It is well known, chromatin accessibility data (e.g. ATAC-seq peaks) profiles all accessible regions in the genome and are composed of different kinds of protein-DNA binding regions. However, most of the current approaches are developed for mainly solving a binary classification task that distinguishes accessible regions from inaccessible regions. Hence, a novel method for constructing a multi-classification task that discriminates different kinds of protein-DNA binding regions is urgently required, which will be helpful for deeply studying the complex protein-DNA binding activities. (ii) Since FCNsignal takes DNA sequences as input and uses base-resolution signals as supervised labels, thereby it is easily expanded to other types of base-resolution signals (e.g. histone modification marks) and even can be transformed into a multi-task framework by integrating multiple types of base-resolution signals (e.g. combining TF-DNA binding signals with H3k27ac signals).

## Materials and methods

### Data preparation

To investigate the overall performance of our proposed method, 53 ChIP-seq datasets for sequence-specific TFs in the HeLa-S3, GM12878, and K562 cell lines, as well as 6 ATAC-seq datasets for the A549, GM12878, HepG2, IMR90, K562, and MCF7 cell lines, were downloaded from the ENCODE project [43] (https://www.encodeproject.org/). To ensure the quality of data, datasets with biological replicates were firstly selected. Secondly, the peaks and signals (*p*-value) were derived from the standard data processing pipelines available from the ENCODE DCC Github (https://github.com/ENCODE-DCC). The peaks that were expanded to 1000bp length and corresponding signals were used as the positive samples, while sequences of the same length at 3000bp upstream of the peaks and corresponding signals were used as negative samples. To alleviate the effect of data bias, the signals were normalized by log_10_(1+signals). The accession list of these datasets is given in Supplementary Table 1.

To test the ability of our proposed method in locating TF-DNA binding regions, the whole chromosome 17 on hg38 was downloaded from UCSC (https://hgdownload.soe.ucsc.edu/downloads.html) and all non-redundant peaks of two TFs (CTCF and YY1) were collected from ReMap2020 [44].

To verify the ability of our proposed method in identifying causal SNPs, several SNP groups with strong linkage disequilibrium (LD) effects were collected. Candidate causal SNPs of myeloma consisting of 10 risk variants at 7p15.3 that alter IRF4 binding [45], pan-autoimmune consisting of seven genetic susceptibility variants at 6p23 that influence the binding of eight TFs [46], and chronic lymphocytic leukemia (CLL) consisting of 27 risk variants at 15q15.1 that disrupt the binding of RELA [47] were obtained from the corresponding literature. Besides, we collected a set of genetic variants from a previous study [37], which contains 29 SNPs associated with breast cancer having strong LD with several SNPs. These SNPs are given in Supplementary Table 2.

### The proposed framework

As shown in Fig 1A, FCNsignal is a symmetrical deep neural network like U-net [48], which takes as input DNA sequences and predicts the base-resolution signals. It is composed of (i) an encoder architecture for down-sampling the inputs and extracting sequence-specific features, (ii) a decoder architecture for up-sampling features and modeling the signal of each base, and (iii) a skip architecture for transferring the position information in the encoder to the high-level features in the decoder. In the following, we will give a detailed description of the three architectures sequentially.

#### The encoder architecture

This part consists of three convolutional blocks in which each block is made up of a convolutional layer, a ReLU layer, a max-pooling layer as well as a dropout layer, a bi-directional GRU (Gated Recurrent Unit) layer [49], and a global average-pooling layer. Specifically, the convolutional block is used to gradually reduce the spatial dimension and encode the sequence-specific features. The bi-directional GRU layer is used to learn the long-term dependencies within the sequence-specific features. The global average-pooling layer is employed to capture the global context of DNA sequences. Generally, the calculation process of this part can be described by formulas 1, 2, and 3.

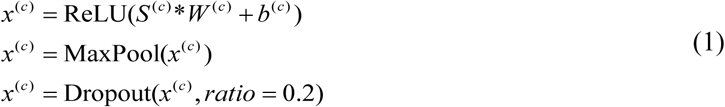

where *S*^(*c*)^, *x*^(*c*)^, *W*^(*c*)^, and *b*^(*c*)^ are input, output, weight matrix, and bias vector of the *c*^th^ convolutional block in the encoder architecture, respectively; * denotes the convolutional operation.

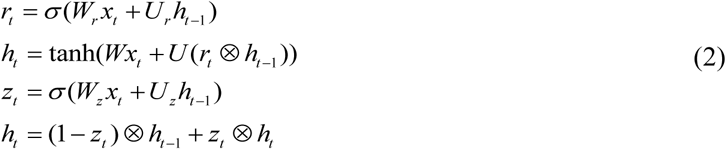

where *r*_*t*_ and *z*_*t*_ are the update gate and the reset gate respectively; *σ* () denotes the logistic sigmoid function; ⊗ denotes the element-wise multiplication operation; *W* and *U* are weight matrices.

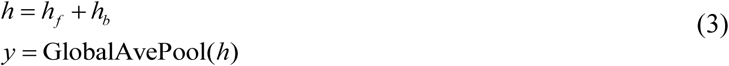

where *h*_*f*_ and *h*_*b*_ are the outputs along with the forward and backward direction.

#### The skip architecture and decoder architecture

This part consists of three skip lines, four upsample layers, and four blending blocks in which each block is composed of a batch normalization (BN) layer, a ReLU layer, and a convolutional layer. Specifically, the skip line is used to combine the position information (‘where’) in the encoder with the discriminating information (‘what’) in the decoder. The upsample layer is used to restore the size of down-sampled features. The blending layer is employed to re-adjust the values of upsampled features. Generally, the calculation process of this part can be described by formula 4.

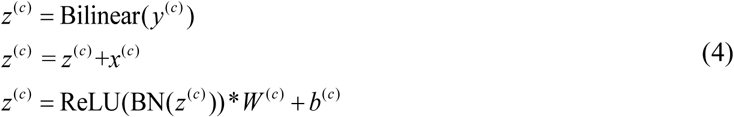

where *W*^(*c*)^ and *b*^(*c*)^ are the weight matrix and bias vector of the *c*^th^ blending block in the decoder architecture respectively; Bilinear(·) is the bilinear interpolation operation; *x*^(*c*)^ is the features from the encoder at the same level. Accordingly, the last output (*z*^(4)^) of this part is regarded as the final prediction of FCNsignal.

The detailed flowchart of FCNsignal is shown in S1 Fig and the architecture settings of FCNsignal can refer to our released codes.

### Model training

Here, we adopted a randomly-split strategy to divide the experimental datasets into training data, validation data, and test data. Specifically, for each dataset, 80% of the peaks with corresponding signals were randomly chosen as the training data, 10% of which were randomly selected as the validation data, while the remaining 20% were used as the test data. The FCNsignal framework transforms TFBSs prediction into a base-resolution regression task, hence the MSE (Mean Squared Error) loss at base-resolution was adopted.

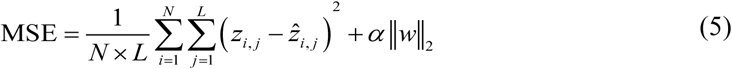

where *N* is the number of samples from the training data; *L* is the length of each sequence; *z*_*i, j*_ and *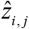* are the predicted and true signal respectively; *α* is a regularization parameter to leverage the trade-off between the goal of fitting and the goal of the generalizability of the trained model; || ||_2_ indicates the L2 norm.

The loss function was optimized by the Adam optimization algorithm [50] with a batch size of 500. The backpropagation algorithm was used for gradient calculating, and exponential decay was applied to the learning rate with a decay rate of 0.9. The weights of our model were initialized by the Glorot initialization method [51]. The learning rate, betas of Adam and regularization parameter were randomly selected from the pre-prepared sets {0.01, 0.001, 0.0001}, {0.9, 0.99, 0.999}, and {0, 0.001} respectively, and the dropout rate was taken from 0.2 or 0.5. These hyperparameters were randomly sampled 15 times, and the validation data were used to select the hyper-parameter set which corresponds to the best performance. Our model was implemented on a single Tesla K40 GPU with 10GB memory using Pytorch.

### Definition of the openness degree of protein-DNA binding

In this study, we defined the openness degree of protein-DNA binding as the number of reads falling into the binding regions, called openness. To achieve this, we firstly collected filtered bam files from ENCODE and then used the ‘multicov’ command in *bedtools* to compute the number of reads falling into each peak. The open degree values for all peaks are further scaled by the log10 function.

### Sequences classification & TFBSs location & motifs prediction

The existing problem is how to use the predicted signals to discriminate binding or non-binding sequences, locate binding regions and predict motifs. In this study, we assumed that, to some extent, the maximum values of signals can reflect the openness degree of TF-DNA binding. To verify this assumption, we computed the Pearson correlation coefficients (Pearsonr) between the openness values and the maximum values of signals. As shown in S11 Fig, the results of four TF datasets confirm that the maximum values of signals are significantly correlated with the openness degree of TF-DNA binding. Therefore, the maximum values of signals were employed to discriminate binding or non-binding sequences as the openness degree of binding sequences is far higher than that of non-binding sequences, and used to locate binding regions as the maximum values are most likely located in the binding regions.

For motifs prediction, binding regions of length 100bp were firstly located by using the position of the maximum values. Secondly, the trained weights of the first convolutional layer were used to select the sub-regions with the highest scores. Thirdly, these selected sub-regions were aligned to compute the corresponding position frequency matrices (PFMs). Finally, TOMTOM [52] was employed to match the PFMs with experimentally validated motifs from standard databases.

### Competing methods and evaluation metrics

To measure the performance of our proposed framework FCNsignal, several existing state-of-the-art methods were used, including MEME [12], STREME [13], DanQ [29], DeepCNN [53], FCNA^*^ [31], FCNA, BPNet [33], DeepEmbed [54], Deopen [55], LSGKM [56] and DeltaSVM [57]. Specifically, MEME discovered TF-DNA binding motifs by searching for repeated, ungapped sequence patterns that occur in the biological sequences. STREME identified ungapped motifs with recurring, fixed-length patterns that are enriched in query sequences or relatively enriched in them compared to control sequences. DanQ predicted TF-DNA binding motifs and prioritized functional SNPs by combining CNN with RNN. DeepCNN, having similar architecture to DeepSea [28], used three convolutional layers to predict TF-DNA binding sites. FCNA^*^ and FCNA shared the same architecture but used different supervised labels, where FCNA^*^ used the base-resolution labels (0/1) which were annotated by using position counting matrices (PCMs) collected from the HOCOMOCO database [58] while FCNA used the base-resolution signals derived from high-throughput experiments. BPNet introduced a deep dilated CNN to predict base-resolution ChIP-nexus binding profiles of pluripotency TFs. DeepEmbed was developed to address the problem of chromatin accessibility prediction via a convolutional Long Short-Term Memory (LSTM) network and *k*-mer embedding. Deopen applied a hybrid deep CNN to learn regulatory sequence code and predict chromatin accessibility at the whole genome level. LSGKM proposed a new version of gkm-SVM for large-scale datasets which offers much better scalability and provides further advanced gapped *k*-mer based kernel functions. As a result, LSGKM achieved considerably higher accuracy than the original gkm-SVM [15]. DeltaSVM presented a new sequence-based computational method to predict the effect of regulatory variation, using a classifier (gkm-SVM) that encodes cell-type-specific regulatory sequence vocabularies.

Since our method involves multiple tasks, several evaluation metrics were adopted, including AUC (Area Under the Receiver Operating Characteristic Curve), PRAUC (Area Under the Precision-Recall Curve), MSE (Mean Squared Error), Pearsonr (Pearson Correlation), and –log_2_(p-value). AUC and PRAUC were used to evaluate the classification performance of FCNsignal. MSE and Pearsonr were used to evaluate the regression performance of FCNsignal. –log_2_(p-value) was used to evaluate the motif prediction performance of FCNsignal.

### Computing SNP scores

Given a specific cell line, we firstly trained FCNsignal with related ATAC-seq data. Secondly, for each SNP, we determined a region of 1000bp around the position of SNP and predicted the signals for the corresponding reference and alteration sequences, respectively. Thirdly, we computed the absolute value of the difference between the two signals at the position of SNP as the SNP score which is equal to 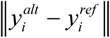, where *i* denotes the position of SNP. Finally, the SNP score was used to identify causal SNPs.

## Availability

The source code of FCNsignal can be found at http://github.com/turningpoint1988/FCNsignal. A web server for FCNsignal is freely available at http://www.imlsb.com:5000/home. The data accession list can be found in S1 Table.

## Supporting information

**S1 Text**. A brief description of applying FCNA to predict motifs on negative sequences.

**S1 Fig**. A detailed flowchart of FCNsignal.

**S2 Fig**. Some examples for showing that FCNsignal can capture the binding signals of TFBSs.

**S3 Fig**. Performance comparison of FCNsignal and the competing methods.

**S4 Fig**. The distribution of the maximum values of signals in the positive and negative samples for CEBPB, CUX1, TCF, and CTCF.

**S5 Fig**. Performance comparison of LSGKM, BPNet, and FCNsignal on the sorted TF ChIP-seq datasets.

**S6 Fig**. Performance of FCNA^*^ on the positive and negative sequences.

**S7 Fig**. Performance of FCNsignal in locating binding regions.

**S8 Fig**. Examples of inputs of arbitrary length for CTCF and YY1.

**S9 Fig**. Pearson correlation coefficients of predicted maximum signals and true maximum signals across 6 ATAC-seq datasets.

**S10 Fig**. The performance of BPNet in identifying causal SNPs from LD groups.

**S11 Fig**. The Pearson correlation coefficients for four TFs including CTCF, YY1, FOSL1, and USF1.

**S1A Table**. The accession list of ChIP-seq datasets from GM12878, K562, and HeLa-S3 cell lines.

**S1B Table**. The accession list of ATAC-seq datasets for A549, GM12878, K562, IMR90, HepG2, and MCF7 cell lines.

**S2 Table**. The SNP list of Autoimmune, Myeloma, and CLL diseases.

**S3 Table**. The motif visualization of three examples (BATF, CTCF, and NFYB).

## Author contributions

**Conceived the basic idea and designed the overall analyses:** D.S. Huang.

**Wrote the manuscript:** Q.H. Zhang.

**Data interpretation and analysis:** S.G. Wang.

**Designed specific experiments:** Q.H. Zhang, Z.H. Chen, and Z. Cui.

**Designed the web server:** Y. He and Z.H. Guo.

